# Phenotypic integration of post-germination traits in *Quercus suber*: morphological development is mediated by acorn mass; leaf physiology by populations’ aridity

**DOI:** 10.1101/2025.01.24.634723

**Authors:** Eduardo Vicente, Marion Carme, Filipe Costa e Silva, Boutheina Stiti, Natalia Vizcaíno-Palomar, Marta Benito Garzón

## Abstract

**Background and Aims:** Assessing intra-specific trait covariation across populations is essential to understand species’ adaptive responses to climatic variation. However, in tree species, this is understudied for early-life stages despite they are more vulnerable to environmental changes and that climatic adaptations can differ between tree ages. In this paper, we studied the integrated phenotype of *Quercus suber* during the months following germination. For that, we studied the covariation of key traits involved in seedlings’ water and C economies along a gradient of aridity at seed origin.

**Methods:** We performed a provenance trial with 157 *Q. suber* seedlings originating from 7 different populations across the species distribution. The seedlings were germinated and grown during 4 months under common conditions. Acorn mass along with 11 above- and below-ground traits involved in water and C use were measured. Their covariation in response to aridity at seed origin was analyzed using structural equation modelling (SEM). The variation of individual traits to increasing aridity and the mediation of acorn mass was also tested.

**Key Results:** Seedlings from arid populations displayed higher leaf evaporative demand coupled with a greater root water uptake capacity. Their leaf physiology also depicted a greater C acquisition capacity, strongly linked to traits conferring drought and heat tolerance. The development of above- and below-ground tissues responded mainly to acorn mass, whereas leaf physiology variations were associated to populations’ aridity.

**Conclusions:** Dry-origin seedlings display a more acquisitive strategy at the whole-plant level compared with seedlings from mesic provenances. This allows a greater water and carbon uptake capacities following germination, which is critical for their survival during their first summer. Leaf physiology adjustments to populations’ climate contrasts with its plasticity observed by other studies addressing juvenile trees, highlighting *Q. suber* varying adaptive strategies at different life stages.

## Introduction

Seedling establishment is one of the most critical steps in the life-cycle of trees and an important bottleneck in forests’ demography (Canham and Murphy, 2016; Ibáñez et al., 2007; Larson et al., 2015; Vanderwel et al., 2013). In Mediterranean ecosystems, this process is quite sensitive to droughts and high temperatures, particularly during the seedlings’ first summer (Martínez-Vilalta and Lloret, 2016; Matías et al., 2011). Nonetheless, in the Mediterranean basin, this can be accentuated in the next decades in a context of more frequent droughts and heatwaves (IPCC, 2022), which can trigger or exacerbate populations’ decay and species’ niche shifts (Fernández-Manjarrés et al., 2018; Gea-Izquierdo et al., 2021; Hartmann et al., 2022). The magnitude of these changes may largely depend on the different adaptive and plastic responses that species present across their distribution range due to different selective pressures between populations (Des Roches et al., 2017; Valladares et al., 2014). For that reason, the study of intra- specific adaptations to varying climate has been a central research field in tree ecology during the last decade. However, most of current knowledge on that matter is based in adult and juvenile trees (Jiménez-Alfaro et al., 2016; Larson and Funk, 2016) despite adaptations and climatic tolerance may differ notably between tree ages (Du et al., 2019; Erlichman et al., 2024; Mašek et al., 2021; Vizcaíno-Palomar et al., 2020), leaving an important knowledge gap in the early stages after germination.

Tree fitness under varying moisture and temperature is mediated by the adjustment of above- and below-ground morpho-functional traits that modulate water and carbon economies (Brodribb et al., 2020; Del Campo et al., 2020; Peguero-Pina et al., 2020). Furthermore, under scarcity or stressful conditions, such as droughts or in arid populations, trees tend to shift their traits to promote conservative resource use strategies (Reich, 2014). For instance, trees in arid populations tend to present adjustments such as reduced stomatal size or density, lower leaf area, or lower specific leaf area (i.e. the mass invested per unit of leaf area, increasing leaf thickness, which promote a lower evaporative demand and higher water use efficiency (Didion-Gency et al., 2021; Mclean et al., 2014; Peguero-Pina et al., 2014). Similarly, at the below-ground level, trees under drier environments tend to present denser root tissue or lower specific root area, which increases their persistence under stressful conditions (De La Riva et al., 2021, 2018; Roumet et al., 2016), as well as deeper or longer roots to ensure access to water resources (Barbeta and Peñuelas, 2016; De La Riva et al., 2018).

Functional traits modulating trees’ resource use often present consistent coordinated responses when exposed to environmental or climatic variation, i.e. trait coordination or phenotypic integration (Messier et al., 2018, 2017). Hence, trees can maintain their fitness status through the synchronized response of multiple traits involving different organs (Kramer-Walter et al., 2016; Metz et al., 2023; Mommer and Weemstra, 2012). For instance, in European *Quercus* species, coordinated responses between specific leaf area (SLA), root-shoot ratio (RS) and photosynthetic rate (A) have been observed in trees under drought treatments, with higher root investment acompassing higher SLA and A (Ramírez-Valiente et al., 2020; Rowland et al., 2023; Solé-Medina et al., 2022). Similarly, Santini et al., (2019) reported that *Pinus halepensis* populations that developed deeper roots consistently increased their spring photosynthetic capacity. Nonetheless, most studies addressing tree responses to climate do so on a single-trait, univariate approach, which provide limited information about their adaptive strategies and can overestimate their responses to future climate (Rowland et al., 2023). In this context, the study of phenotypic integration has been proposed as a more insightful methodology to understand trees adaptive processes (Messier et al., 2017; Poorter et al., 2014). Particularly, the use of trait integration networks or structural equation models (SEM) is incipient in trait-based ecology, allowing to identify key trait groups or specific trait combinations that are promoted by climatic selection (Benavides et al., 2021; Santini et al., 2019; Torres-Ruiz et al., 2019; Vicente et al., 2022).

Understanding species’ adaptive strategies and their potential responses to future climate is of special importance for wide-range species whose climatic niche may undergo important shifts in the following decades. In this context, *Quercus suber* L. (cork oak) is a relevant species (Benito Garzón et al., 2008). It spans along the Mediterranean basin, with an important presence in the west Iberian Peninsula. Its populations cover a wide climatic variability, from the more mesic and cool conditions in the south of France, to the arid and warm climate in the Maghreb region (Durrant et al., 2016). This species is adapted to drought and has a great resistance to fire (Burrows and Chisnall, 2016; Pausas, 2015, 1997), which confer it a great ecological and reforestation interest in a climate change context. In this line, previous studies have deemed populations from the southern and more arid distribution as interesting genetic resource based on their higher fitness in common gardens and due to genetic differentiation in traits related to drought resistance (Morillas et al., 2023; Ramírez-Valiente et al., 2017, 2014, 2010). However, there is a lack of information about whether these trends are valid concerning these species’ early life stages after germination. For instance, Benito Garzón et al., (2024) reported significant earlier germination times in acorns from southern populations, which may expose seedlings to a higher frost mortality risk if these acorns are planted in colder-than- origin locations. Nevertheless, seedlings adaptive strategies after germination, and particularly regarding their phenotypic integration, is yet to be explored.

In this paper, we carry out a study of the phenotypic integration of *Q. suber* post- germination morpho-functional traits in response to populations’ aridity. For that, we study the variation of different fitness-related traits in 4 month-old seedlings raised in common conditions and coming from seven populations covering the species’ climatic niche. We put our focus in seed properties (mass) as well as in above- and below-ground seedling traits regulating water and carbon economies. Our goal is to determine (1) which adaptive-trait configurations are promoted by increasing aridity, (2) how these traits co- vary between them in response to increasing aridity along the species range, and (3) which are the main traits and functions mediating such responses. We expect that the seedlings from the more arid populations will show morphological adjustments that allow a greater water uptake (e.g. greater development of fine roots and main root length). We also expect that acorn mass will mediate these morphological adjustments. In addition, we hypothesize that this greater water uptake capacity in dry-origin seedlings will be consistently linked to adjustments aimed to minimize water loss by evapotranspiration (e.g. reductions in leaf area, SLA and stomatal size or density).

## Material and methods

### Acorn sampling, germination, and nursery

Acorns were harvested during autumn in 2022 from 7 populations across the distribution range of *Q. suber* (Fig. 1, table 1). All acorns were collected at the same ripening stage (i.e. brown pericarp and without radicle emergence). No acorns from the ground were picked up, and only those that fell from the trees when shaking the branches were collected. In every population, at least 10 trees were sampled with a minimum of 10 acorns per tree. Distance between mother trees was always greater than 30m. After collection, acorns were sent immediately to the laboratory, arriving within 1 – 5 days. Sealed bags with humid cloths and cold gels were used to keep humidity and cold during transport.

**Figure 1:**
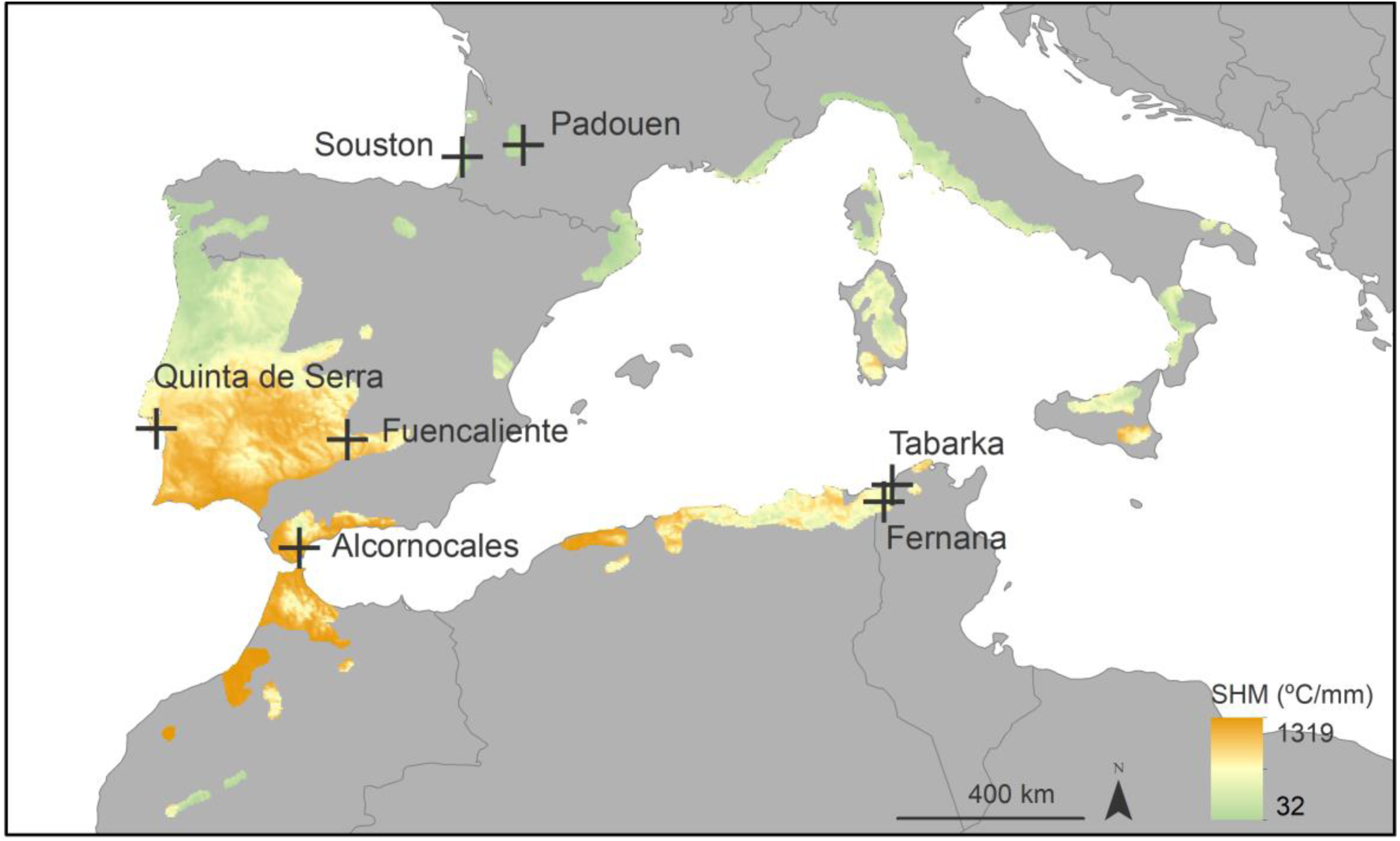
Map of *Q. suber* distribution and sampled populations. Color gradient depicts the current (2021) summer heat moisture index along *Q. suber* geographic distribution. The *Q. suber* distribution map was downloaded from the database compilation by Caudullo et al., (2017).

**Table 1.** Description of the sampled populations.

| Population | Country | Latitude | Longitude | Elevation<br>(m.a.s.l.) | SHM<br>(°C.mm<sup>-1</sup>) | Seedlings | Mothers |
| --- | --- | --- | --- | --- | --- | --- | --- |
| Souston | France | 43.894 | -1.346 | 23 | 49.43 | 24 | 9 |
| Padouen | France | 44.092 | 0.297 | 101 | 71.93 | 13 | 6 |
| Fuencaliente | Spain | 38.421 | -4.313 | 748 | 211.79 | 31 | 10 |
| Alcornocales | Spain | 36.302 | -5.431 | 66 | 464.65 | 23 | 7 |
| Quinta de Serra | Portugal | 38.488 | -9.027 | 109 | 275.92 | 17 | 6 |
| Tabarka | Tunisia | 36.952 | 8.855 | 43 | 257.02 | 28 | 8 |
| Fernana | Tunisia | 36.654 | 8.609 | 421 | 225.65 | 21 | 7 |

Acorns were checked for viability and sown immediately after receiving them. Viability was assessed by a floating test, submerging them in water during 5 minutes. Floating acorns were subsequently discarded. Then, viable acorns were sown in nursery trays (4x5 cells) with cell dimensions of 6 x 6 x 8 cm each, filled with universal substrate (NPK 8- 2-7, dry organic matter 80%, conductivity 30 mS/m, pH 6.5, water retention 820 ml/L). The individual fresh mass of every acorn (AM) was noted before sowing. In total, 294 acorns from 57 mother trees were sown. The acorns were sown by half burying them in the soil with the apex facing up. In every tray, each line corresponded to a family (mother tree) of the same population. The trays were then placed in a climatic chamber (Snijder LABS, micro clima-series) at 20°C with a constant 75% relative humidity, 300 µmol m^−2^ s^−1^ light intensity, and a photoperiod of 13/11 light/dark hours. During the dark phases the temperature was lowered by 5 °C in all chambers. Trays were manually watered twice a week with distilled water until saturation.

The trays remained for germination and early seedling growth in the chamber during approximately 2 months. Once they had emerged, seedlings were manually watered once a week. After that period, the seedlings were placed in bigger pots of 18 cm depth to avoid constrains in root development. However, these bigger pots allowed limited vertical space for shoot growth in the climatic chambers. For that reason, the seedlings were moved to a lab nursery with a temperature variation of 22-18 °C, and natural daylight cycle. They remained in the nursery for another 2 months with the same watering treatment.

### Functional traits measurements

After 2 months in the nursery, we measured several seedlings’ morphological and functional traits related to resource use economy. The number of sampled seedlings is indicated in table 1.

First, while seedlings were still in the pots, we estimated leaf chlorophyll (Chl), flavonol (Flv) and anthocyanin (Ant) contents nondestructively using a MPC-100 multi-pigmentmeter (ADC BioScientific Ltd, UK) on a fully developed and exposed leaf. Then, we measured seedling height (H, cm), number of leaves, number of branches, and the leaf area (cm^2^) of a representative leaf (i.e. fully developed, exposed, and from the mid-upper stem section) using the Petiole Pro smartphone application (Petiole, LTD. https://www.petiolepro.com/). Then, this metric was used to estimate total leaf area (LA, cm^2^) by multiplying it by the seedling’s leaf number. Stomatal length (SL, μm, used as surrogate of stomatal size thereafter), width (SW, μm), surface (SS, μm^2^), and density (SD, n stomata.mm^-2^) were further measured on a representative leaf. For that, we obtained nail varnish imprints from a mid-leaf location separated from the central vein. The imprints were photographed using a Leica DM 2500 microscope (x40 magnification) equipped with a Leica MC190 HD camera (Leica Microsystems GmbH, 300ppp). The resulting pictures were then analyzed using image J software. Finally, all the aboveground biomass was dried at 60 °C during 3 days to obtain the stem and leaves dry weights. The dry mass of the leaf used to calculate LA was weighted separately and used to estimate specific leaf area (SLA, m^2^. kg ^-1^).

In addition, we cleaned and scanned the seedlings’ root system using an EPSON Perfection V850 Pro scanner (EPSON America, Inc., USA). The main root and the fine roots were processed separately. The obtained images were analyzed with the RhizoVision Explorer software v.2.0.3 (Seethepalli et al., 2021) to calculate main and fine roots’ total length (MR_L and FR_L, cm), area (MR_A and FR_A, cm^2^) and volume (MR_V and FR_V, cm^3^). After that, we also measured the main root’s collar using a digital caliper (RC, mm). Then, both main and fine roots were dried at 60 °C during 3 days to obtain their dry biomass. We further used these dry weights to estimate main- and fine roots specific root length (SRL_MR and SRL_FR, m.kg^-1^), specific root area (SRA_MR and SRA_FR, m^2^.kg^-1^), and root tissue density (RTD_MR and RTD_FR, kg.m^-3^). Finally, root-shoot ratio (RS, unitless) was calculated dividing the whole root dry weight by the total aboveground dry biomass.

### Climate data

To characterize each population climate, we used several variables related to temperature and precipitation from the Climate DT database (https://www.ibbr.cnr.it//climate-dt/).

These variables were spring maximum temperature (Sp_Tmax, °C), minimum temperature (Tmin, °C), mean spring precipitation (MSP, mm), precipitation during the growing season (GSP, April-October, mm), temperature seasonality (T_seasonality), and the summer heat moisture index (SHM, °C mm^-1^). SHM is defined as MWMT/(MSP/1000) (Marchi et al., 2024), where MWMT is the Mean Warmest Month Temperature (°C) and MSP the Mean Summer Precipitation (mm). We chose these variables because aridity, as well as high and low temperatures during spring and summer are the main abiotic filters for *Q. suber* seedlings’ for early establishment (Ramírez-Valiente et al., 2022). Each climatic variable was averaged for the period 1901-1960, as this period is a better representative of the climate shaping potential adaptations in the mother trees rather than today’s values.

### Statistical analysis

#### Phenotypic integration along the climatic gradient

To analyze the phenotypic integration along the seed provenances’ climatic gradient we used a structural equation model (SEM) approach. To avoid overfitting the model with many traits representing the same function, or that may present a great collinearity between them, we performed a previous PCA for the above- and below-ground traits separately (Figs. S1-2). We did that in order to select the suit of traits that best represented, from an ecological point of view, the morphology, resource-use economy and physiology of the sampled populations. As aboveground traits, we finally selected LA, SLA, Chl, Flv and SL. Concerning belowground traits, we selected MR_L, FR_A, SRA_MR and SRA_FR. Despite their ecological importance, we did not include H and RS. H was highly correlated with LA, while in the case of RS, we wanted to avoid using ratios due to their potential linear dependence with other traits included in the SEM model.

Likewise, we performed another PCA using the selected traits and the climatic variables downloaded from ClimateDT, to check which of these were better associated to the functional traits’ variability (Fig. S3). All the climatic variables contributed mainly to the first PCA dimension, indicating a strong correlation between them in our dataset. The aridity index, SHM, was the variable with the highest score in the PCA, therefore we chose it as the potential driver of traits adjustment for the analysis.

Then, we defined 7 latent variables: one to describe the climatic gradient across our populations, “Aridity”, defined by the SHM index; and another one to account for the effects of acorn properties, “Acorn mass”, defined by AM. The other 5 latent variables described the functional properties associated to the selected traits. Specifically, these 5 latent variables were: “Development”, to describe the growth of morphological tissues related to resource use and acquisition, defined by LA, MR_L and FR_A; “Leaf display”, defined by SLA; “Root display”, defined by SRA_MR and SRA_FR; “Acquisitive leaf physiology”, defined by Chl and SL; and “Leaf protection pigments”, defined by Flv. It must be noted that, initially, our conceptual model split Development in two different latent variables for aboveground and belowground traits. However, we finally decided to integrate them in a single one due to a strong correlation between LA and FR_A that prevented model convergence. A summary of the selected traits, their associated functions and assigned latent variables is provided in table 2.

**Table 2.** Description of the measured functional traits, related functions and associated latent variables.

| Trait |  | Description | Function | Latent variable |
| --- | --- | --- | --- | --- |
| *Seedling development - growth traits* |  |  |  |  |
| **FR_A** | Fine roots' area | Fine roots' total surface | Water uptake, nutrients absorption | Development |
| **MR_L** | Main root lenght | Tap root's lenght | Rooting depth, water uptake | Development |
| **LA** | Total leaf area | Sedlings' total foliar surface | Evapotranspiration, water demand | Development |
| **H** | Height | Seedlings' shoot lenght | Growth, competition for light | - |
| *Leaf and roots' economics tratis* |  |  |  |  |
| **SLA** | Specific leaf area | Leaf surface / leaf dry mass (of a representative leaf) | Fast leaf development, schlerophylly | Leaf display |
| **SRA_FR** | Specific root area (fine roots) | Fine roots' total surface / Fine roots' dry mass | Fast root development, root persistence | Root display |
| **SRA_MR** | Specific root area (main root) | Main roots' total surface / Main roots' dry mass | Fast root development, root persistence | Root display |
| *Allocation traits* |  |  |  |  |
| **RS** | Root-Shoot ratio | Belowground dry mass / Aboveground dry mass | Investment in belowground or aboveground tissue | - |
| *Stomatal traits* |  |  |  |  |
| **SL** | Stomatal size (lenght) | Mean stomatal pore lenght | Transpiration, carbon uptake | Acquisitive leaf physiology |
| *Leaf pigments* |  |  |  |  |
| **Chl** | Chlorophyll content | Ratio T850nm / T720nm | Carbon uptake | Acquisitive leaf physiology |
| **Flv** | Flavonol content | Ratio fluorescence F660nm / F325nm | Leaf protection against light, temperature, drought | Leaf protection pigments |
| *Seed traits* |  |  |  |  |
| **AM** | Acorn mass | Acorn fresh weight | Cotyledon reserves | Acorn mass |

The final SEM included direct regressions between Aridity and the rest of the latent variables, to account for adjustments directly related to the populations’ climate. Similarly, to account for adjustments mediated by AM, the model included direct regressions between Acorn mass and the seedlings’ latent variables. Finally, the model did not include direct regressions between the seedlings’ latent variables and we only allowed free covariation between them (Fig. S4).

The SEM model was fitted using the R package *lavaan* (Rosseel, 2012). The goodness of fit was evaluated using the X^2^ statistics, which for SEMs indicate a good fit when p > 0.05. Nonetheless, large sample sizes tend to produce low p regardless of the model fitness, thus, providing more fitness statistics is strongly recommended when fitting SEMs (Beaujean, 2014). For that reason, we also checked the comparative fit index (CFI) which indicates a good fit when is higher than 0.90 (Bentler, 1990); the root mean square of approximation (RMSEA), depicting a good fit when values are below 0.06-0.08 (Awang, 2012); and the standardized root mean square residuals (SMRS) which indicates a good fit when is lower than 0.09 (Hu and Bentler, 1990). All traits and variables included in the SEM where centered and scaled.

#### Models for individual traits

In addition to the phenotypic integration approach, we analyzed each trait variation along the aridity gradient individually, in order to have a detailed insight about their response to the seed provenances’ climatic cline and the influence of acorn mass, while also accounting for the structure of the dataset (seedlings coming from the same mother, and several mothers per population). To that purpose, we fitted linear mixed models (LMM) for the same traits included in the SEM analysis. In addition, these individual models were also fitted for seedlings’ height (H) and root-shoot ratio (RS).

The models took the following general form:

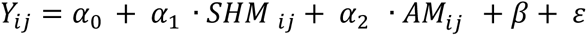

Where Yij is the trait measured for the ith seedling of the jth population. SHM is the aridity index of the j population, AM the acorn fresh weight that produced the ith seedling of the jth population, β are the random effects and ε the residuals. The random effects included mother tree nested to population. These models were fitted using the *lmer()* function of the R package *lme4* (Bates et al., 2015). The variance explained by the models was assessed calculating the marginal (R^2^M, i.e. considering only the fixed effects) and conditional (R^2^C, i.e. considering both fixed and random effects) variances using the function *r.squaredGLMM()* of the R package *MuMIn* (Barton, 2020).

## Results

### Phenotypic integration along the gradient

The goodness of fit indices obtained in the SEM model (df = 27, X^2^ = 45.09, p = 0.016) depicted an overall good fit, with CFI = 0.949, SRMR = 0.062 and RMSEA = 0.069 (Fig. 2, table 3).

**Figure 2:**
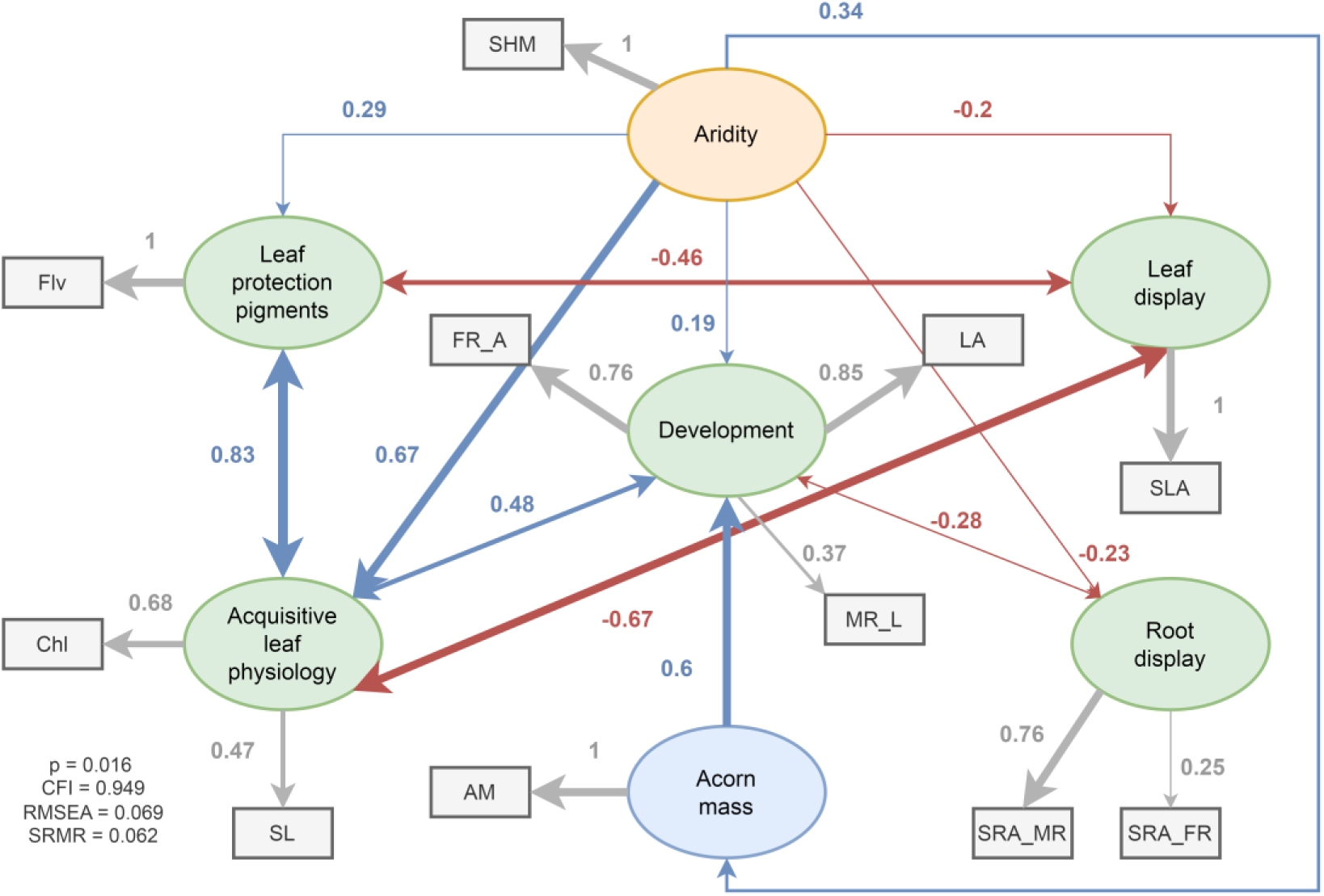
Structural equation model showing latent (ellipses), observed (rectangles) variables and the relationships between them. The relationships between variables are represented with standardized path coefficients. Only significant paths (p<0.05) are represented. Blue arrows depict positive relationships, red ones depict negative relationships, and grey ones depict the factor loading of each observed variable to the corresponding latent variable. For the observed variables, SHM stands for Summer Heat Moisture index; AM for acorn fresh weight, LA for total leaf area; MR_L for main root length; FR_A for fine roots’ area; SL for stomatal size; Chl for chlorophyll content; SLA for specific leaf area, Flv for flavonol content, SRA_MR and SRA_FR for specific root area of main and fine roots’, respectively.

**Table 3.** Parameter estimates of the Structural Equation Model. =∼ refers to the contributions to the latent variables; ∼ to regressions; ∼∼ to covariances between latent variables. SHM stands for Summer Heat Moisture index; AM, acorn fresh weight; LA, total leaf area; MR_L, main root length; FR_A, fine roots’ area; SL, stomatal size; Chl, chlorophyll content; SLA, specific leaf area, Flv, flavonol content, SRA_MR and SRA_FR, specific root area of main and fine roots.

| Latent variable |  |  | est | SE | Z | p |
| --- | --- | --- | --- | --- | --- | --- |
| Aridity | =~ | SHM | 1 |  |  |  |
| Development | =~ | LA | 1 |  |  |  |
| Development | =~ | FR_A | 0.905 | 0.104 | 8.718 | 0 |
| Development | =~ | MR_L | 0.448 | 0.109 | 4.106 | 0 |
| Acquisitive leaf physiology | =~ | Chl | 1 |  |  |  |
| Acquisitive leaf physiology | =~ | SL | 0.727 | 0.134 | 5.435 | 0 |
| Leaf protection pigments | =~ | Flv | 1 |  |  |  |
| Acorn mass | =~ | AM | 1 |  |  |  |
| Leaf display | =~ | SLA | 1 |  |  |  |
| Root display | =~ | SRA_MR | 1 |  |  |  |
| Root display | =~ | SRA_FR | 0.364 | 0.294 | 1.238 | 0.216 |
| Regressions |  |  |  |  |  |  |
| **Development** | ~ | **Aridity** | **0.166** | **0.069** | **2.416** | **0.016** |
| **Development** | ~ | **Acorn mass** | **0.517** | **0.071** | **7.299** | **<0.001** |
| Acorn mass | ~ | **Aridity** | **0.342** | **0.08** | **4.281** | **<0.001** |
| **Acquisitive leaf physiology** | ~ | **Aridity** | **0.44** | **0.074** | **5.909** | **<0.001** |
| Acquisitive leaf physiology | ~ | Acorn mass | -0.134 | 0.071 | -1.888 | 0.059 |
| **Leaf protection pigments** | ~ | **Aridity** | **0.293** | **0.087** | **3.35** | **0.001** |
| Leaf protection pigments | ~ | Acorn mass | -0.086 | 0.087 | -0.991 | 0.322 |
| **Leaf display** | ~ | **Aridity** | **-0.202** | **0.09** | **-2.26** | **0.024** |
| Leaf display | ~ | Acorn mass | 0.078 | 0.089 | 0.875 | 0.382 |
| **Root display** | ~ | **Aridity** | **-0.166** | **0.084** | **-1.982** | **0.047** |
| Root display | ~ | Acorn mass | -0.087 | 0.083 | -1.056 | 0.291 |
| Covariations |  |  |  |  |  |  |
| **Development** | ~~ | **Acquisitive leaf physiology** | **0.146** | **0.051** | **2.865** | **0.004** |
| Development | ~~ | Leaf protection pigments | 0.000 | 0.059 | 0.008 | 0.994 |
| Development | ~~ | Leaf display | 0.016 | 0.061 | 0.257 | 0.797 |
| **Development** | ~~ | **Root display** | **-0.115** | **0.058** | **-1.988** | **0.047** |
| **Acquisitive leaf physiology** | ~~ | Leaf protection pigments | **0.390** | **0.072** | **5.375** | **<0.001** |
| **Acquisitive leaf physiology** | ~~ | **Leaf display** | **-0.323** | **0.071** | **-4.562** | **<0.001** |
| Acquisitive leaf physiology | ~~ | Root display | -0.010 | 0.059 | -0.164 | 0.870 |
| **Leaf protection pigments** | ~~ | **Leaf display** | **-0.423** | **0.085** | **-4.955** | **<0.001** |
| Leaf protection pigments | ~~ | Root display | -0.028 | 0.072 | -0.381 | 0.703 |
| Leaf display | ~~ | Root display | 0.103 | 0.075 | 1.379 | 0.168 |

The results point out that, along the studied climate gradient, aridity increases were related with heavier acorn mass (path coefficient = 0.34), higher development of morphological tissues related to resource acquisition (path coefficient = 0.19), a more acquisitive leaf physiology (path coefficient = 0.67) and greater concentration of leaf protection pigments (path coefficient = 0.29). Conversely, greater aridity was associated with significant reductions in leaf- and root display (path coefficient = -0.2 and -0.23, respectively). Acorn mass only showed a significant influence on Development, which increased when Acorn mass was heavier (path coefficient = 0.6).

Among the seedling functions defined by the latent variables, Acquisitive leaf physiology showed a strong positive covariation with Leaf protection pigments (path coefficient = 0.83) and Development (path coefficient = 0.48), while it had a negative covariance with Leaf display (path coefficient = -0.67). Leaf display also showed a negative relationship with Leaf protection pigments (path coefficient = -0.46), while Root display showed a similar covariation with Development (path coefficient = -0.28).

Additionally, it is worth noting that fine roots’ area (FR_A) and total leaf area (LA) were the functional traits that contributed most to the latent variable Development (factor loadings of 0.76 and 0.85, respectively), while the role of main root length (MR_L) was relatively small (factor loading = 0.37). A similar situation was observed for the latent variable of Root display, where main root’s specific root area (SRA_MR) was the major contributor (factor loading = 0.76) over fine roots’ SRA (SRA_FR; factor loading = 0.25).

### Individual trait models results

The models fitted for individual traits (Table S1) showed that all the considered morphological traits related to seedling development were significantly modulated by acorn mass, but not by SHM (Fig. 3, table S1). Specifically, higher acorn mass significantly increased seedling height (t = 5, R^2^M = 0.17, R^2^C = 0.45), total leaf area (t = 5.67, R^2^M = 0.29, R^2^C = 0.46), main root length (t = 3.76, R^2^M = 0.1, R^2^C = 0.13) and fine roots’ area (t = 5.34, R^2^M = 0.31, R^2^C = 0.44), while none of them showed a significant response to the aridity index SHM.

**Figure 3:**
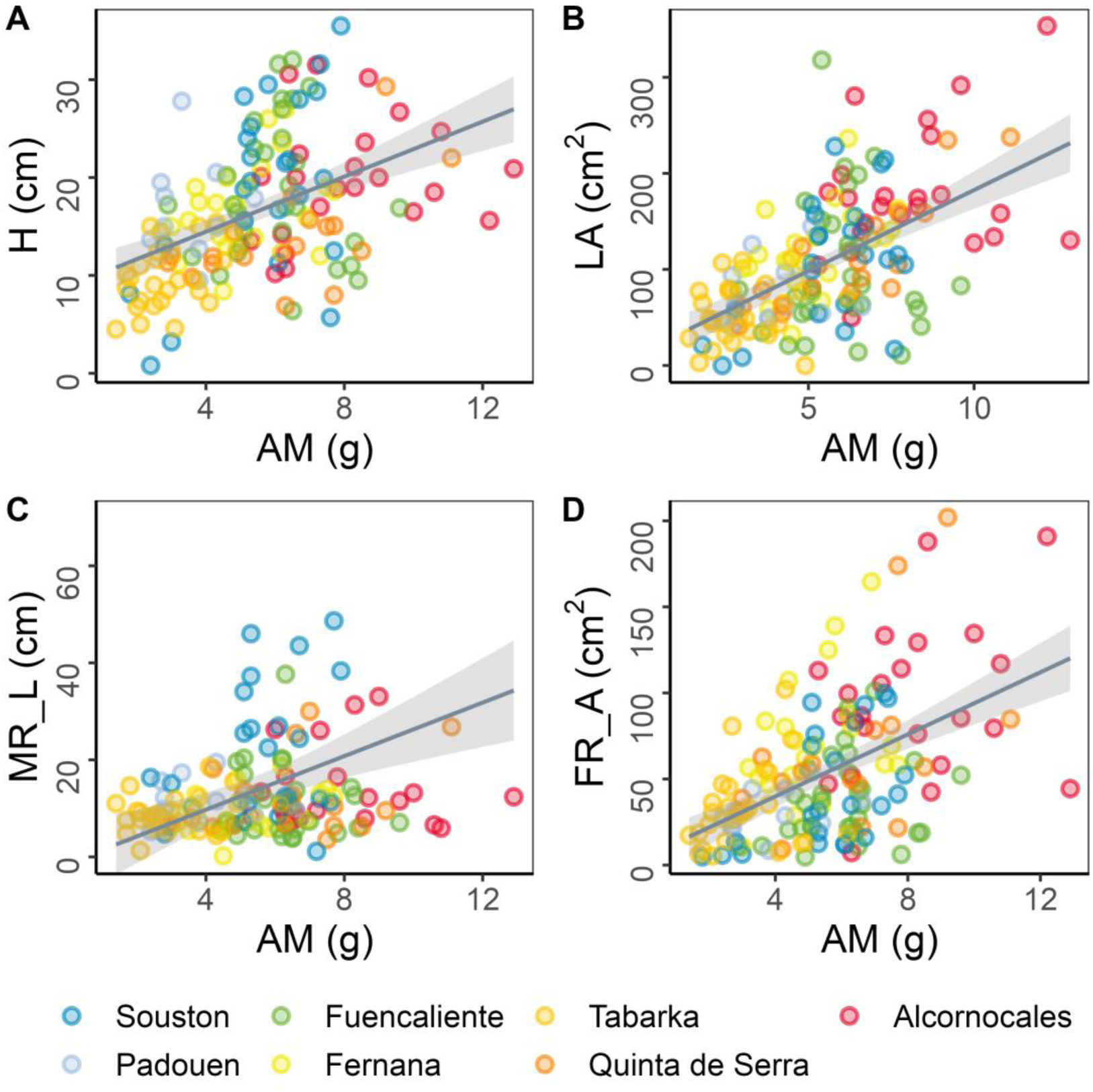
Response of A, seedling height (H); B, total leaf area (LA); C, main root length (MR_L); and D, fine roots’ area (FR_A) to acorn mass (AM) variation. Solid trend lines depict significant effects.

On the contrary, the traits related to leaf physiology showed significant increases with higher SHM, whilst acorn fresh weight had no significant effect in their response (Fig. 4, Table S1). Particularly, higher SHM increased Chl (t = 1.96, R^2^M = 0.18, R^2^C = 0.61) and SL (t = 2.22, R^2^M = 0.14, R^2^C = 0.25). The models also depicted a significant reduction of the main root’ SRA when aridity increased (t = -2.24, R^2^M = 0.06, R^2^C = 0.13, Fig. 4, Table S1). Conversely, neither SRA_FR, SLA, Flv, nor RS ratio (Table S1) showed significant responses to the variation of SHM and AM when analyzed with LMM. The model for acorn fresh weight also depicted no significant influence of aridity on this trait (Table S1, Fig. S5).

**Figure 4:**
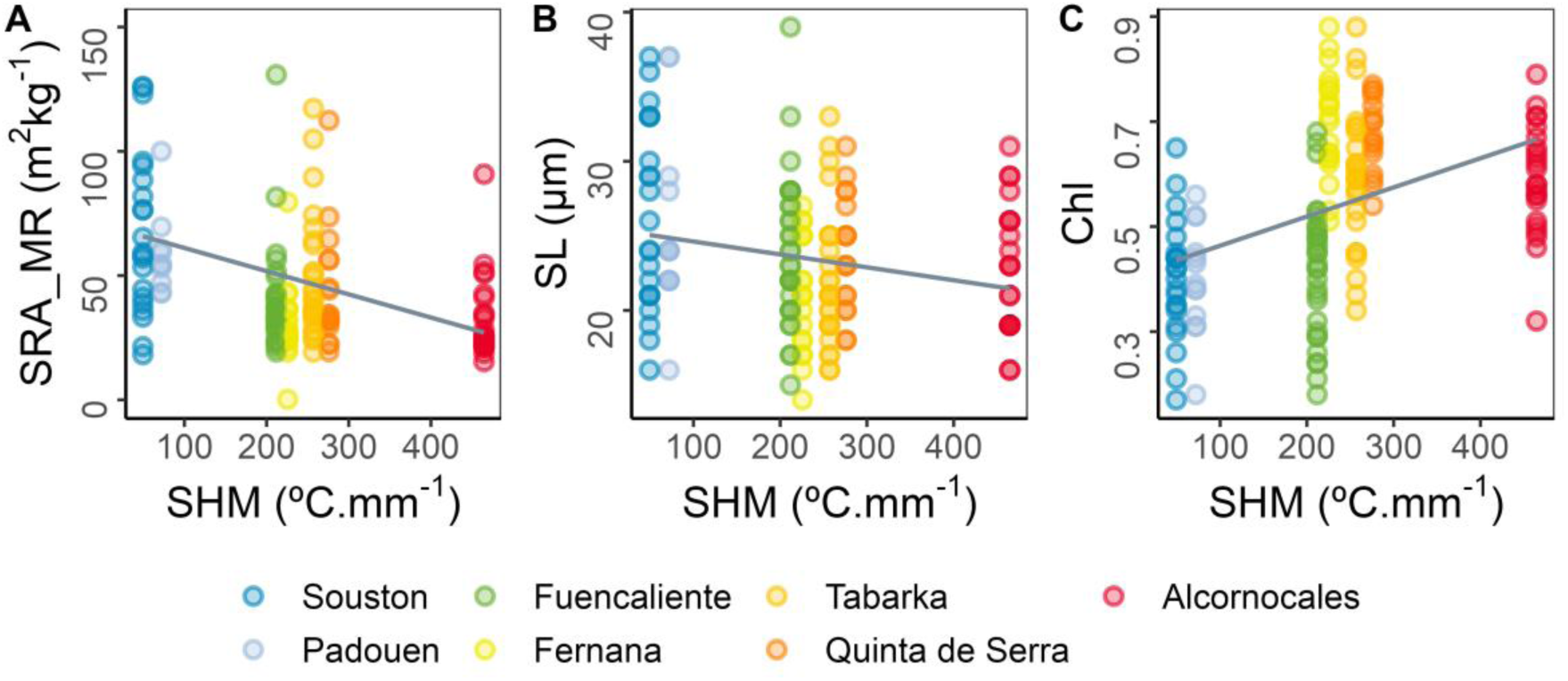
Response along the aridity (SHM) gradient of: A, specific root area of the main root (SRA_MR); B, stomatal size (SL); C, chlorophyll content (SL). Solid trend lines depict significant effects.

## Discussion

In this paper, we studied the post-germination integrated phenotype of *Q. suber* seedlings coming from seven populations scattered along an aridity gradient covering the species’ niche. These seedlings were germinated and grown during 4 months under common conditions, which allowed to identify selection patterns associated to populations’ climate. Seedlings from more arid populations displayed a greater development of tissues involved in water acquisition and use. This was, at the same time, tightly linked with leaf physiological traits involved in C acquisition, and the latter were associated to greater leaf protection against desiccation and high temperatures. In addition, our results showed that the development of above- and below-ground tissues was determined mainly by acorn mass, whilst leaf physiology variations were strongly linked to aridity at seed origin.

### Seedlings’ phenotypic integration in response to aridity at seed origin

The phenotypic integration analysis depicted that, as populations’ aridity increased, seedlings showed a greater development of above- and below-ground morphological traits involved in water use. Specifically, this response was mainly based on a synchronized increment of total leaf area (LA) and fine roots area (FR_A). The tight coordination between these traits depicts an adjustment at root level to compensate for leaves’ water demand, which may be of particular importance for seedlings’ fitness under dry conditions. Indeed, seedlings with higher LA display a greater evaporative demand, and hence they are also exposed to a higher dehydration and mortality risk under drought conditions (Matías et al., 2019b; Morillas et al., 2023). However, an increase of FR_A implies an important increment in the soil volume explored for hydric resources. Furthermore, water uptake by the root system is mainly due to the activity of the fine roots (Padilla and Pugnaire, 2007; Weemstra et al., 2020). Thus, FR_A increases can notably enhance water uptake and promote a well hydrated and physiological status (Otieno et al., 2006; Ramírez-Valiente et al., 2018). Accordingly, greater investments in roots are commonly reported in literature as an adaptive adjustment of Mediterranean oaks inhabiting dry and drought-prone environments (Lázaro-González et al., 2023) including *Q. suber* (Matías et al., 2019a; Morcillo et al., 2020; Ramírez-Valiente et al., 2022, 2018). These adjustments are often depicted an increase in the root-shoot ratio (RS), however, this contrasts with our results, as RS remained constant along the studied provenances. Nonetheless, Morillas et al., (2023) also observed similar RS ratios between *Q. suber* seedlings from different provenances several months after germination, while also reporting greater development of roots and leaves in dry-origin ones. Together with our results, this highlights the important link between roots and leaves development in the post-germination phase of *Q. suber* seedlings.

Our results also point out that leaf physiology traits varied strongly in response to populations’ aridity. Particularly, seedlings from more arid populations showed a more acquisitive leaf physiology, based on greater chlorophyll content (Chl) and stomatal size (SL). Higher Chl promotes an enhanced photosynthetic capacity, and greater SL has been associated to increased stomatal conductance in *Q. suber* (Prats et al., 2019). In addition, these physiological adjustments were tightly coupled with a greater concentration of leaf flavonols (Flv) and with more sclerophylous leaves (i.e. lower SLA). Leaf flavonols provide protection against abiotic stress, mainly dissipating the impacts of high temperatures and excessive UV radiation, and its accumulation is often increased in plants inhabiting drought-prone environments (Laoué et al., 2022; Nakabayashi et al., 2014; Sharma et al., 2019). Similarly low SLA is a known adaptive response of Mediterranean plants growing under dry conditions and frequently reported in *Quercus* species (Leiva and Fernández-Alés, 1998; Ramírez-Valiente et al., 2020; Ramírez-Valiente et al., 2010, 2009). This adjustment is often linked to greater leaf thickness and enhances protection against excessive dehydration, further allowing to extend photosynthetic activity during drought conditions (Didion-Gency et al., 2021; Ramírez-Valiente et al., 2010). Taken together, these results point out that seedlings from more arid populations displayed an acquisitive and drought-tolerant leaf physiology after germination, allowing a greater carbon acquisition beyond favorable periods of water availability and mild temperatures.

In addition, our results point out an important positive covariation between the seedlings’ acquisitive leaf physiology and the development of leaves and roots. This highlights an overall acquisitive strategy at plant level in dry-origin seedlings. In this line, it must be noted that their morphological traits’ display can be expensive in terms of carbon economy (De La Riva et al., 2021; Eissenstat et al., 2000), and although C demands for initial seedling enlargement are usually covered by cotyledon reserves (Mechergui et al., 2021), their higher assimilation capacity may be aimed to compensate for growth demands. Nonetheless, an acquisitive strategy during the months following germination can be essential for seedling survival during their first summer, especially in the more arid populations. Indeed, traits promoting enhanced water use and carbon acquisition are sometimes observed in tree populations inhabiting seasonal or unpredictable climates (Hajek et al., 2014; Kiorapostolou et al., 2020). This allows to maximize C assimilation and growth when water availability is not limiting, compensating for scarcity periods such as drought spells or long winters. For instance, greater water transport capacity and leaf surface have been reported in *Fagus sylvatica* populations at the species’ southern distribution limit, more prone to seasonal droughts and heatwaves (Vicente et al., 2022). Similarly, arid populations of *Quercus oleiodes* have been observed to display higher leaf N content and C assimilation capacity than mesic ones (Ramírez-Valiente et al., 2017). These adjustments allow to maintain an overall positive C balance over time, and to relocate resources in key organs or to increase reserves (Eilmann et al., 2009). Indeed, in Mediterranean species, higher C reserves are related to a greater resilience facing severe disturbances such as extreme droughts (Lloret et al., 2018; Martínez-Vilalta et al., 2016; Morcillo et al., 2022) as they can promote a quicker growth recovery, the formation of new conductive tissue, or resprouting (Barbeta and Peñuelas, 2016).

Indeed, resprouting capacity is a key trait underlying *Q. suber* resilience and regeneration capacity after extreme events such as fire or droughts. *Q. suber* trees can quickly recover from stem buds, effectively protected by the species bark (Burrows and Chisnall, 2016; Pausas, 2015). However, in the case of young seedlings, the stems are very thin and more prone to die, thus their resprouting capacity relies on buds from the stem base or the main root (Pausas, 1997). In this line, our results from both the trait integration and individual trait models point out a higher investment in the main root of dry-origin seedlings, depicted by a significant increase of the main root specific root area (SRA). This suggests a denser root tissue (De La Riva et al., 2021), which increases its resistance and seedlings’ recovery chances following eventual dieback during their first summer.

### Leaf physiology is linked to provenance climate, but morphology and growth are controlled by acorn mass

Both our phenotypic integration and individual trait variation approaches highlight that SL and Chl were the traits most strongly modulated by populations’ aridity. Nonetheless, this result contrasts with common garden experiments with oak species that did not detect provenance differentiation in stomatal, photosynthetic, or water use efficiency traits, which highlights that trait variability is rather due to plasticity than to adaptation. For instance, for *Q. suber*, Ramírez-Valiente et al., (2009) found no difference in water use efficiency of saplings from different populations after several years growing in common environments. Similar results have been reported for *Quercus petraea*, including stomatal size and density (Torres-Ruiz et al., 2019). In addition, chlorophyll content is a trait that can vary greatly depending on temperature, light, and moisture availability (Aranda et al., 2008; Fang and Xiong, 2015; Lázaro-González et al., 2023). Nonetheless, the divergence we observed in these traits between the wet and arid populations suggest their importance during the first stages after germination. In this line, it must be noted that *Quercus* species’ acorns are recalcitrant seeds that do not tolerate desiccation (Joët et al., 2016, 2013), thus, germination occurs under mild water availability conditions. Then, the observed acquisitive leaf physiology in seedlings from the more arid populations might be tuned with these mesic conditions following germination, which would allow to maximize carbon uptake and allocation in key organs.

Conversely, growth and morphological traits, especially the development of total leaf area and fine roots’ mass, were mainly mediated by acorn fresh weight. Indeed, there is broad evidence in literature depicting that seed mass is directly related to early seedling growth and survival (Mechergui et al., 2021; Wright and Westoby, 1999; Yi et al., 2015), which has been observed in many Mediterranean *Quercus* species such as *Q. ilex, Q. faginea* and *Q. suber* (Lázaro-González et al., 2023; Morillas et al., 2023; Ramírez-Valiente et al., 2009). This greater development in seedlings from heavier acorns is explained by a larger embryo as well as by higher cotyledon reserves (Jurado and Westoby, 1992; Milberg and Lamont, 1997; Yi et al., 2015), which promote greater growth rates and seedling size. Higher shoot and root growth confers seedlings a greater capacity to access and mobilize resources (Villar-Salvador et al., 2012), allowing them to maintain their activity and fitness during periods of elevated evapotranspiration such as summer drought. In addition, larger acorns are also reported to store a greater water content (Amimi et al., 2023), which reduces their desiccation risk between shedding and germination (Joët et al., 2016).

In that line, broad-range studies with Mediterranean *Quercus* species often report acorn size variations along aridity gradients, with larger acorns observed in dryer and warmer provenances at the species’ southern limits (Benito Garzón et al., 2024; Morillas et al., 2023; Ramírez-Valiente et al., 2009). Accordingly, our results depict a divergent acorn mass between the mesic and arid populations in our gradient, suggesting that bigger acorns may be selected in the most arid end of our gradient to indirectly increase seedling development and survival chances. However, our analyses yielded an overall weak effect of populations’ aridity on acorn mass due to the small size of the Tunisian ones, suggesting that other factors rather than climate had also an influence on this trait in our dataset. A potential cause, on one hand, may be on the environmental conditions experienced by mother trees during acorn ripening, as too dry or too warm conditions in that stage can significantly reduce the number and size of seeds (Caignard et al., 2021; Fernández-Pascual et al., 2019). Indeed, this effect has been reported in the closely related *Q. ilex* when environmental conditions in September are drier than usual (Alejano et al., 2011). On the other hand, this could also be related to a different genetic lineage in the Tunisian trees. Particularly, Vanhove et al., (2021) observed that *Q. suber* populations from the east Mediterranean basin, including those in Tunisia, belong to a differentiated lineage from the ones in the Iberian Peninsula and the Atlantic coast.

### Perspectives and limitations

Overall, this study highlights that, during the first months following germination, and under mild conditions, *Q. suber* seedlings from arid provenances display a more resource-acquisitive strategy at plant level compared to seedlings from mesic ones. This strategy was based on a higher development of fine roots tightly coupled with a greater leaf area and shoot length, depicting an enhanced water uptake capacity to compensate leaves water demand. This was acompassed by a leaf physiology allowing a higher carbon acquisition and gas exchange capacities, based on a greater stomatal size and chlorophyll content. In addition, the more acquisitive leaf physiology of dry-origin seedlings was linked to lower SLA and higher content of protection pigments, depicting a greater capacity to extend carbon assimilation under drought conditions. This trait configuration would confer seedlings from arid populations a greater capacity to access and mobilize soil hydric resources, as well as to maximize carbon assimilation during water availability periods after germination, which can be advantageous for their survival and resilience facing their first summer. Therefore, the integrated phenotype of seedlings from arid populations may confer them a greater capacity to maintain their early fitness in a climate change context. Thus, more arid *Q. suber* populations could be considered as an interesting seed source for future reforestation or regeneration programs concerning this species, although higher temperatures can notably hasten their germination phenology (Vicente and Benito Garzón, 2024), increasing hence frost risks during regeneration.

However, it must be noted that the development of morphological traits was determined mainly by acorn mass, highlighting that an important component of the seedlings’ fitness and their potential response to climate change is mediated by this trait. In this regard, the acorn mass sampled in the Tunisian populations was low, which may be concerning taking into account that populations at the southern distribution limit potentially need a greater change in allelic frequencies to adapt to climate change. This result suggest that conservation efforts should also focus on populations from the southern limit of this species. Nonetheless, acorn development can vary greatly depending on the conditions experience by mother trees during ripening (Alejano et al., 2011; Fernández-Pascual et al., 2019), and our study was performed with acorns from a single year. Thus, further studies are needed to assess annual variations of acorn mass along the species distribution, as well as their potential effects in seedlings early fitness.

The variability in between the leaf physiology sensitivity to populations’ aridity observed in our study during the post-germination stage and its absence in saplings (Ramírez-Valiente et al., 2009) demonstrate the changing climate sensitivity of *Q. suber* at different life stages. While these variations have been predicted by mathematical models (Erlichman et al. 2024), they are rarely verified through empirical data. Such differences in climate sensitivity across the tree’s lifespan should be considered in assisted migration and gene flow programs that aim to optimize seed source selection in response to climate change (Chakraborty et al., 2024). Moreover, our findings reinforce the need to shift breeding objectives from focusing on single traits to promoting integrated phenotypes that enhance drought tolerance (Climent et al., 2024).

## Acknowledgments

We are grateful to Julien Goullier (Le Liege Gascon) for his assistance during acorn collection ; Chloée Jean for her exhaustive technical support during acorn collection, seedling monitoring and functional traits’ measurements; Inès Moreau for her help during root scanning and image processing; and Soizig Le-Stradic for her advice on roots’ processing and root traits measurements.

## Authors’ contribution

EV and MBG designed the study. All authors participated in acorn collection. EV, MC and MBG performed the lab work and seedlings’ measurements. EV analyzed the data and leaded the writing of the manuscript. All authors contributed critically to the writing of the manuscript.

## Data and materials accessibility statement

The data supporting the findings of this study will be publicly available in the Zenodo repository upon manuscript acceptance.

