## Supplementary figures and tables for "Phenotypic integration of post-germination traits in *Quercus suber*: morphological development is mediated by acorn mass; leaf physiology by populations’ aridity"

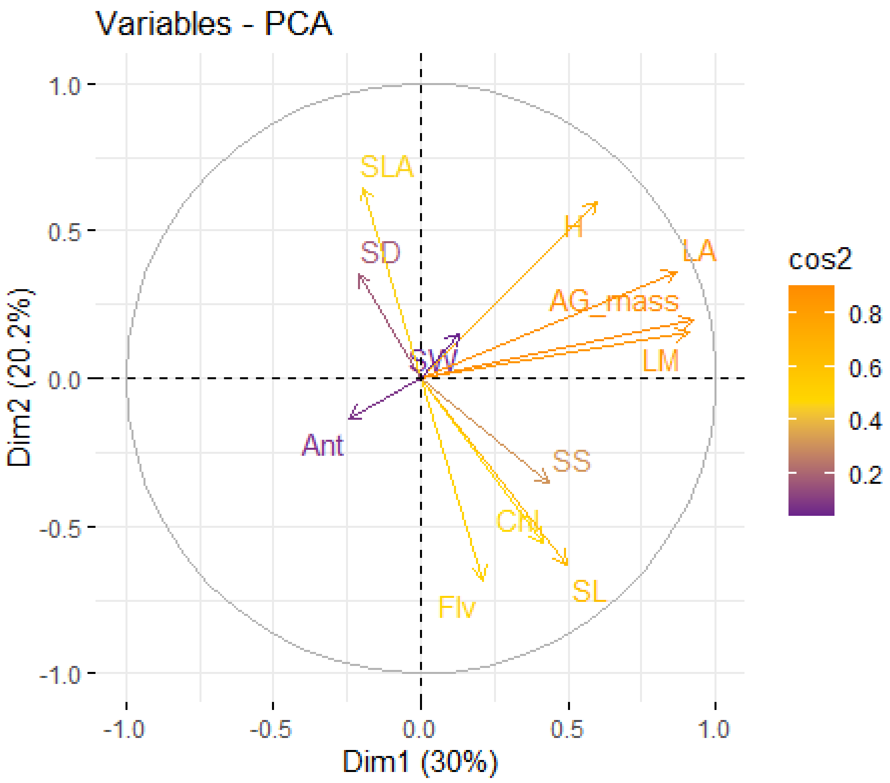


**Figure S1.** PCA of the measured aboveground traits. SLA stands for specific leaf area; H, seedling height; LA, total leaf area; AG_mass, total aboveground dry mass; LM, total leaves' dry mass; SS, stomatal surface; Chl, chlorophyll content; SL, stomatal size; SD, stomatal density; SW, stomatal width; Flv; flavonols content; Ant, Anthocyanin content.

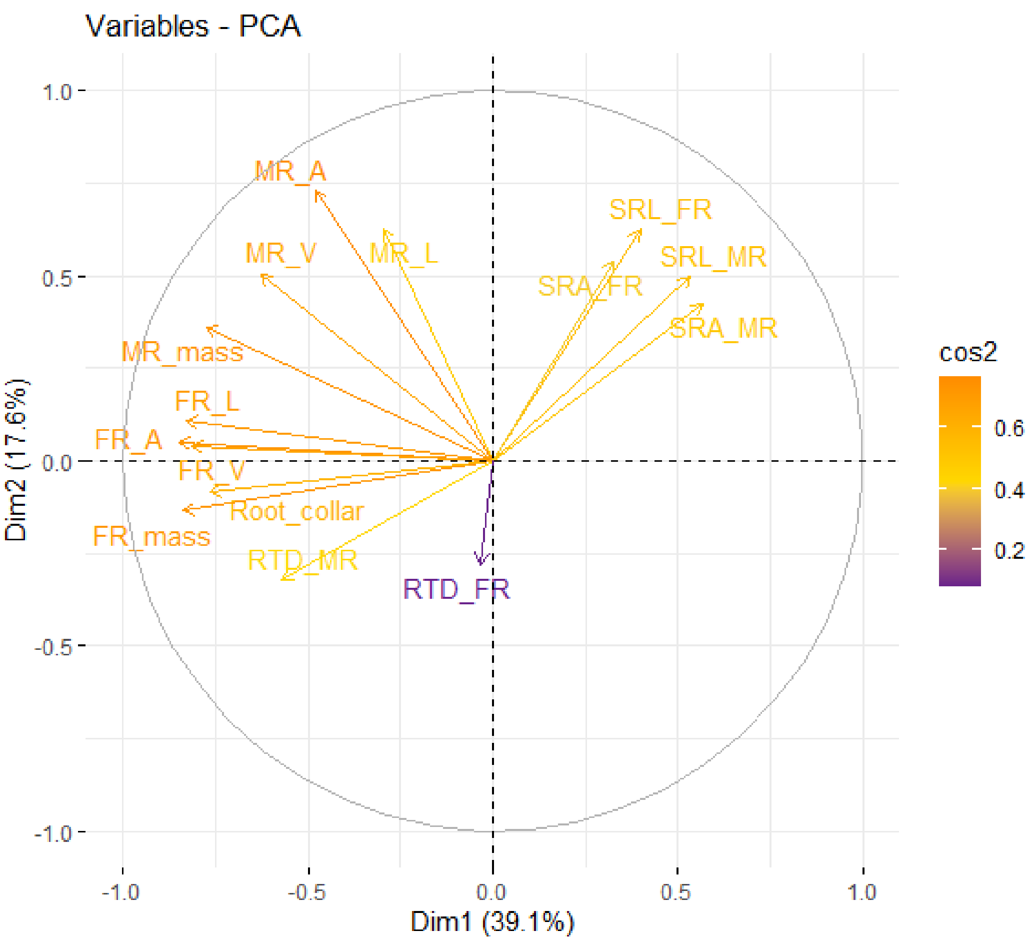


**Figure S2.** PCA of the measured belowground traits. For main roots (MR) and fine roots (FR), L stands for length; A for area; V for volume; and mass for dry biomass. SRL stands for specific root length; SRA, specific root area; RTD, root tissue density.

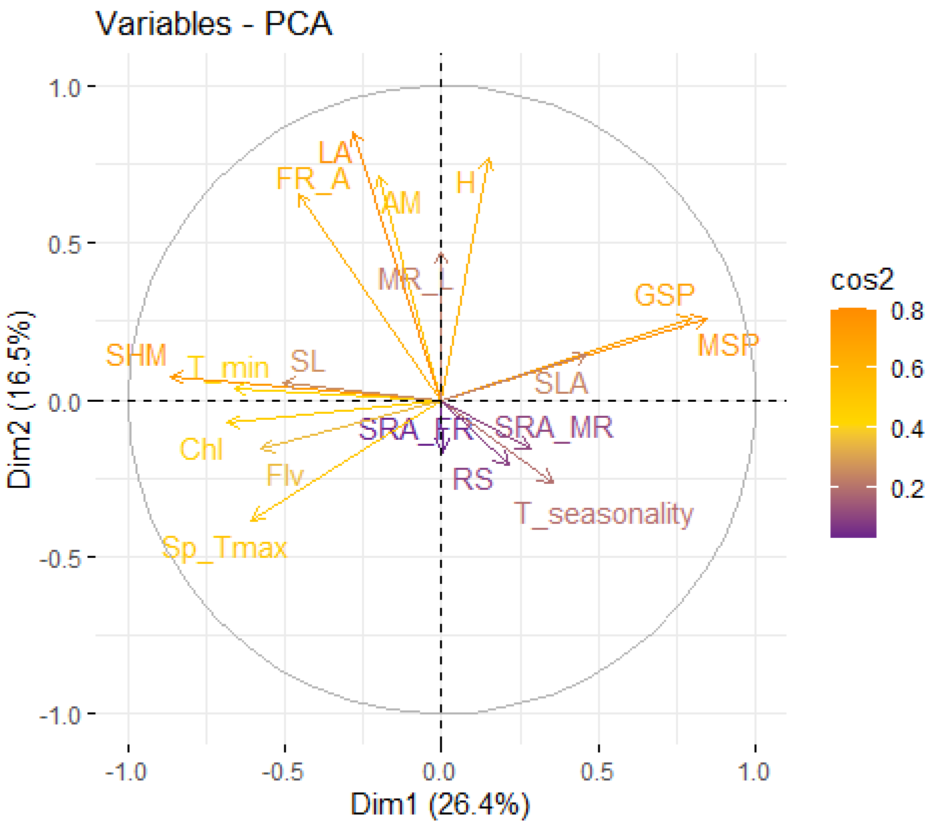


**Figure S3.** PCA of the considered traits with the climatic variables. SHM stands for Summer heat moisture index; MSP, mean summer precipitation; GSP, growing season precipitation; Tmin, minimum temperature; Sp_Tmax, maximum spring temperature, T_seasonality; temperature seasonality; RS, root-shoot ratio; H, seedling height; AM, acorn fresh weight; LA , leaf area; MR_L, main root length; FR_A, fine roots' area; SL, stomatal size; Chl, chlorophyll content; SLA, specific leaf area; Flv, flavonol content; SRA_MR and SRA_FR, specific root area of main and fine roots, respectively.

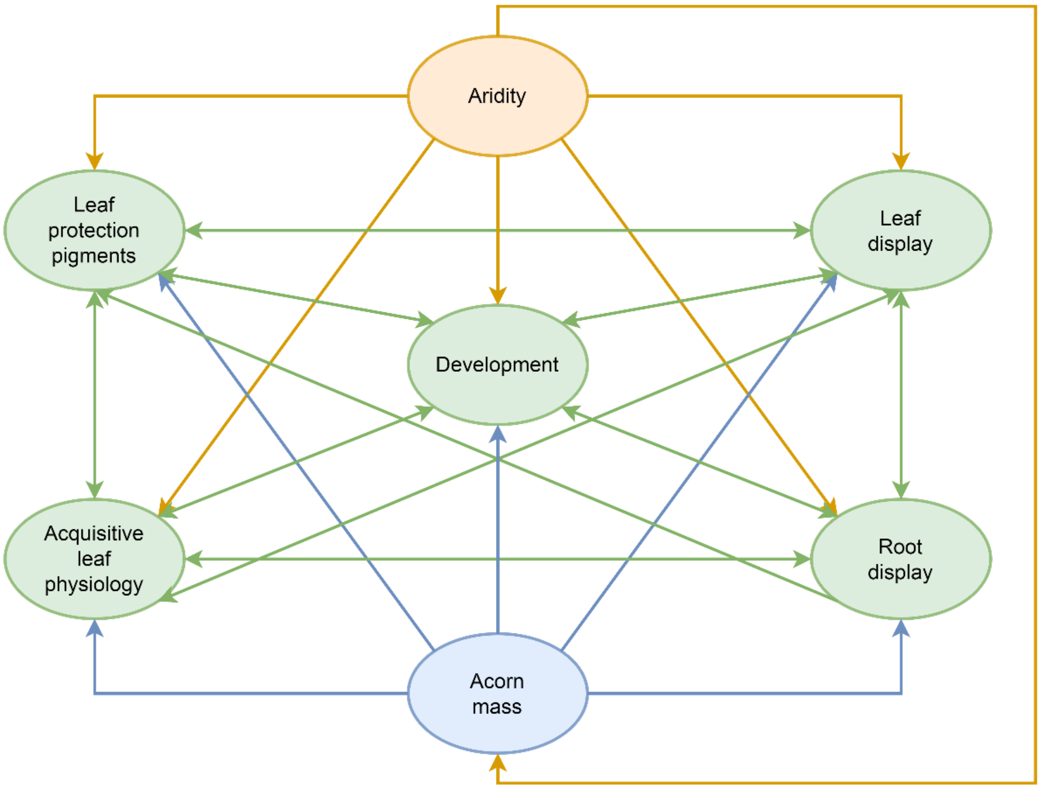


**Figure S4.** Conceptual representation of the SEM model, depicting relationships between the aridity gradient, seed, and seedling traits. Relationships involving aridity are represented in orange color; relationships involving acorn mass in blue color; relationships between seedling traits in green color.

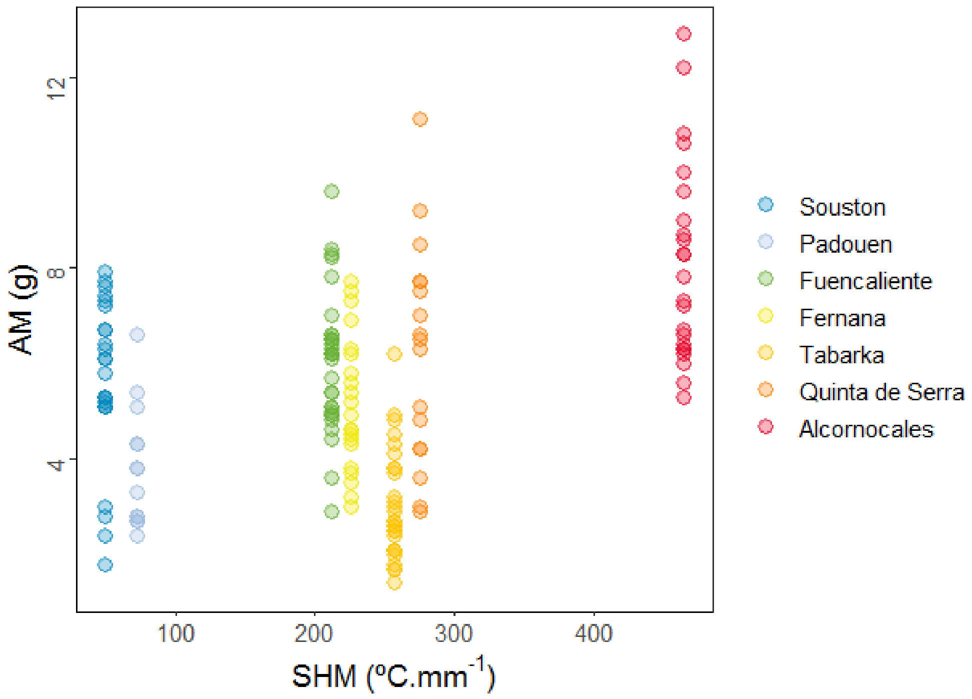


**Figure S5.** Distribution of acorn mass along the aridity gradient.

**Supplementary tables.**

**Table S1.** Summary of the individual traits' LMM. AM, acorn fresh weight; H, seedling shoot height; LA, total leaf area; MR_L, main root length; FR_A, fine roots' area; SLA, specific leaf area; SRA_MR and SRA_FR, specific root area of main and fine roots; SL, stomatal size; Chl, chlorophyll content; Flv, flavonol content; RS, root-shoot ratio; SHM, summer heat moisture index; DBH, mother trees' diameter at breast height.

| **Trait** | | **Fixed effects** | | **Random effects** | | **Model fitness** | |
|---|---|---|---|---|---|---|---|
| | | SHM | AM | | Variance | $R^2$M | $R^2$C |
| **AM** | est | 0.34 | x | Prov | 0.39 | 0.11 | 0.65 |
| | SE | 0.24 | x | Prov/mother | 0.19 | | |
| | t | 1.42 | x | | | | |
| **H** | est | -0.19 | **0.43** | Prov | 0.05 | 0.17 | 0.45 |
| | SE | 0.12 | **0.08** | Prov/mother | 0.21 | | |
| | t | -1.52 | **5.00** | | | | |
| **LA** | est | 0.15 | **0.46** | Prov | 0.04 | 0.29 | 0.46 |
| | SE | 0.11 | **0.08** | Prov/mother | 0.12 | | |
| | t | 1.35 | **5.67** | | | | |
| **MR_L** | est | -0.06 | **0.34** | Prov | 0.02 | 0.1 | 0.13 |
| | SE | 0.10 | **0.09** | Prov/mother | 0 | | |
| | t | -0.62 | **3.76** | | | | |
| **FR_A** | est | 0.23 | **0.45** | Prov | 0.09 | 0.31 | 0.44 |
| | SE | 0.13 | **0.08** | Prov/mother | 0.04 | | |
| | t | 1.69 | **5.34** | | | | |
| **SLA** | est | -0.13 | -0.16 | Prov | 0.32 | 0.05 | 0.33 |
| | SE | 0.22 | 0.09 | Prov/mother | 0 | | |
| | t | -0.58 | -1.65 | | | | |
| **SRA_MR** | est | **-0.23** | -0.06 | Prov | 0.02 | 0.06 | 0.13 |
| | SE | **0.10** | 0.09 | Prov/mother | 0.04 | | |
| | t | **-2.24** | -0.71 | | | | |
| **SRA_FR** | est | 0.01 | -0.02 | Prov | 0 | 0.01 | 0.01 |
| | SE | 0.09 | 0.09 | Prov/mother | 0 | | |
| | t | 0.10 | -0.26 | | | | |
| **SL** | est | **0.34** | 0.10 | Prov | 0.12 | 0.14 | 0.25 |
| | SE | **0.15** | 0.09 | Prov/mother | 0 | | |
| | t | **2.22** | 1.10 | | | | |
| **Chl** | est | **0.48** | -0.08 | Prov | 0.41 | 0.18 | 0.61 |
| | SE | **0.24** | 0.08 | Prov/mother | 0.07 | | |
| | t | **1.96** | -1.02 | | | | |
| **Flv** | est | 0.31 | 0.01 | Prov | 0.55 | 0.08 | 0.54 |
| | SE | 0.28 | 0.09 | Prov/mother | 0.04 | | |
| | t | 1.11 | 0.08 | | | | |
| **RS** | est | -0.08 | 0.10 | Prov | 0.04 | 0.01 | 0.21 |
| | SE | 0.12 | 0.10 | Prov/mother | 0.16 | | |
| | t | -0.67 | 1.00 | | | | |

**Table S2**. Mean values ± SD of the measured trait values averaged by provenance. AM, acorn fresh weight; H, seedling shoot height; LA, total leaf area; MR_L, main root length; FR_A, fine roots' area; SLA, specific leaf area; SRA_MR and SRA_FR, specific root area of main and fine roots; SL, stomatal size; Chl, chlorophyll content; Flv, flavonol content; RS, root-shoot ratio.

| | **Souston** | **Padouen** | **Fuencaliente** | **Fernana** | **Tabarka** | **Quinta de Serra** | **Alcornocales** |
|---|---|---|---|---|---|---|---|
| **AM (g)** | 5.65 ± 1.7 | 3.85 ± 1.2 | 6.05 ± 1.4 | 5.16 ± 1.4 | 3.04 ± 1.1 | 6.23 ± 2.3 | 8.12 ± 2.1 |
| **H (cm)** | 19.16 ± 9.14 | 16.77 ± 4.47 | 18.60 ± 6.99 | 17.35 ± 4.66 | 10.05 ± 3.23 | 14.01 ± 5.34 | 19.92 ± 6.03 |
| **LA ($cm^2$)** | 108.09 ± 65.39 | 71.55 ± 33.48 | 100.09 ± 70.93 | 121.02 ± 44.38 | 54.85 ± 29.8 | 111.71 ± 58.58 | 178.22 ± 67.53 |
| **SLA ($m^2$ $kg^{-1}$)** | 296.56 ± 105.58 | 354.33 ± 104.37 | 340.33 ± 132.28 | 253.04 ± 61.73 | 202.30 ± 98.91 | 199.16 ± 43.55 | 284.01 ± 104.92 |
| **Chl** | 0.40 ± 0.11 | 0.40 ± 0.1 | 0.42 ± 0.13 | 0.70 ± 0.1 | 0.60 ± 0.14 | 0.67 ± 0.07 | 0.59 ± 0.11 |
| **Flv** | 0.25 ± 0.12 | 0.24 ± 0.08 | 0.33 ± 0.1 | 0.39 ± 0.12 | 0.42 ± 0.18 | 0.63 ± 0.17 | 0.34 ± 0.11 |
| **SL (µm)** | 0.138 ± 0.013 | 0.128 ± 0.007 | 0.143 ± 0.015 | 0.147 ± 0.013 | 0.146 ± 0.011 | 0.151 ± 0.011 | 0.149 ± 0.012 |
| **RS** | 1.20 ± 0.83 | 0.94 ± 0.41 | 1.49 ± 1.03 | 1.04 ± 0.35 | 1.16 ± 0.62 | 0.96 ± 0.51 | 1.01 ± 0.36 |
| **MR_L (cm)** | 22.08 ± 14.13 | 11.75 ± 4.36 | 10.95 ± 7.19 | 8.73 ± 4.26 | 8.79 ± 3.68 | 11.97 ± 8.19 | 23.60 ± 45.87 |
| **FR_A ($cm^2$)** | 43.42 ± 33.53 | 29.87 ± 13.29 | 37.19 ± 24.41 | 73.22 ± 36.61 | 32.68 ± 23.37 | 65.94 ± 50.74 | 93.64 ± 44.22 |
| **SRA_MR ($m^2$ $kg^{-1}$)** | 76.00 ± 51.6 | 58.44 ± 16.72 | 53.16 ± 75.11 | 30.34 ± 14.52 | 47.76 ± 24.30 | 43.68 ± 23.44 | 33.18 ± 17.04 |
| **SRA_FR ($m^2$ $kg^{-1}$)** | 434.44 ± 146.30 | 492.90 ± 110.32 | 540.85 ± 216.91 | 439.88 ± 90.52 | 479.39 ± 167.28 | 629.55 ± 918.47 | 434.72 ± 111.81 |
